# 3D printed titanium anodized effects on human gingival fibroblasts response and bacterial colonization: a dual approach

**DOI:** 10.64898/2026.03.11.711067

**Authors:** Lanig Lefort, Sébastien Gilles, Susan Chamorro-Rodriguez, Marie-Laurence Giorgi, Stéphane Petit, Audrey Asselin, Christophe Beloin, Benjamin Fournier, Marie-Joséphine Crenn

## Abstract

Mucointegration is as important as osseointegration to ensure the survival of implant-supported prosthesis. Indeed, effective soft tissue integration (STI) prevents the appearance of complication through bacterial dissemination. To optimize STI, electrochemical anodization can be used to nanostructure the trans-gingival part of the prosthetic component. Moreover, Selective Laser Melting (SLM) is a new 3D-manufacturing technique that enables the production of customized implant-supported prosthesis with complex geometry.

**Objective:** The aim of this study is to evaluate the effect of a SLM manufactured and anodized Ti6Al4V surface on the behaviour of both, human gingival fibroblasts and oral bacteria.

**Method:** SLM-Ti6Al4V discs were polished and anodized with defined parameters to obtain nanotubes (NTs) with specific morphology. Surface characterization was assessed through surface topography and wettability. Human gingival Fibroblasts were cultured, and cell morphology was observed by SEM at day 7. Proliferation, viability (day 1,4,7) and adhesion (6 h and 36 h) were analyzed. Then immunofluorescence and RT-qPCR were used to detect the distribution and the gene expression of vinculin at 48 h. An early colonizer (Streptococcus gordonii) was used for a parallel evaluation of bacteriological adhesion.

**Results:** SLM-ANO-Ti6Al4V showed similar performances in terms of cytotoxicity, compared with a machined and polished titanium surface currently used in clinics. Interestingly, cell adhesion was enhanced on anodized SLM surfaces, with a difference in the distribution of focal adhesion plaques in HGFs, while biofilm formation of S. gordonii was not affected by anodization.

**Significance:** SLM anodized surface showed promising ability to promote STI while controlling bacterial adhesion.

## Introduction

Implant-supported prostheses (IPS) represent the treatment of choice for replacing missing teeth, with a survival rate of the implant level at 96.4% after 10 years [1]. However, survival doesn’t mean no complication. Indeed, clinical follow-up studies have shown that only 66% of patients remain free from adverse outcomes after 5 years [2].

One of the most common complications that can affect the long-term success rate of ISPs is peri-implantitis, the most severe form of peri-implant disease. This biological complication is characterised by inflammation of the peri-implant soft tissues and progressive bone loss. This pathology is a major health concern, with an estimated incidence of 21.7% [3], which increases over time [4].

The development of peri-implantitis can be explained by a number of factors, but one of the primary causative agent is believed to be poor plaque control [5]. Since the abutment (mostly made of grade 5 titanium) is an individual transmucosal structure that occupies most of the transmucosal region [6], the current approach to preventing plaque accumulation is to control its shape, and to polish its surface to minimize the roughness of its transgingival portion and thus avoid bacterial adhesion. This is achieved by creating an arithmetic average surface roughness (Ra) below 0.2 µm, which is defined as the threshold below which bacterial colonization becomes negligible [7].

However, the aforementioned “mirror-polished” surface probably influence human gingival cells behavior and could be part of the explanation for the histological difference between the connective tissues around the trangingival and the gingiva around natural teeth [6]. Indeed, collagen fibers do not adhere to the surface of the transgingival component and are organized in parallel around it. Combined with the more limited presence of cells, this result in an inferior transmucosal seal considered to promote the rapid progression of peri-implantitis compared to periodontitis [8].

Then, the main challenge lies in conceiving a surface that promotes gingival cell adhesion, also known as “muco integration”, while simultaneously preventing bacterial colonization.

In order to enhance muco integration, various surface modification strategies have been explored, of which nanostructured titanium surfaces appear to be the most promising [9,10]. The application of nanoscale surface structuring, employing nanotemplates, nanopores, or nanotube arrays, can beneficially alter surface properties to support the attachment and function of human oral cells [11]. A range of techniques have been developed to achieve such nanostructures [12], but electrochemical anodization is distinguished by its simplicity, reproducibility, and cost-effectiveness [9]. When applied under well-controlled conditions, anodization can generate titanium nanotubes (NTs) with adjustable surface characteristics. By modifying the surface topography, chemistry, and wettability, these nanotubular structures have the capacity to influence the behaviour of human cells [13,14], including gingival fibroblasts and keratinocytes residing in peri-implant soft tissues. Numerous *in vitro* studies have reported that NTs alone can enhance the proliferation and adhesion of gingival fibroblasts [15,16]. A recent *in vivo* study conducted on canines reported that titanium-based abutments anodized with nano-sized NTs increased connective tissue length and perpendicular collagen fibres in beagle dogs after a healing period of four weeks, in comparison with machined abutments [17]. Furthermore, NTs have the capacity to function as reservoirs for drugs or bioactive molecules, which can be utilised to inhibit bacterial growth [18] or to enhance the cellular environment through the delivery of growth factors or functional proteins[19,20]. Thus, despite the variability of NT characteristics across studies, titanium nanotubes emerge as a highly promising approach for promoting favourable gingival tissue responses.

The majority of studies conducted on nanostructured titanium have focused on machined or turned titanium alloys. However, it is noteworthy that the abutment may now be produced through the utilisation of additive manufacturing techniques.

Selective Laser Melting (SLM) is a particularly promising approach within this field, offering greater design flexibility and the possibility of patient-specific geometries when compared with conventional milling methods. [21]. Nevertheless, the distinctive microstructure of SLM-fabricated titanium components affects how anodization modifies the surface and consequently influences the resulting surface properties. In recent research, it was demonstrated that, despite the impact of the SLM microstructure on the distribution of nanotubes (NTs), it remains feasible to produce anodized SLM-Ti6Al4V surfaces (that have been previously polished) with promising NTs properties to enhance collagen adsorption [22].

To date, only a few *in vitro* studies have assessed the effect of TiO_2_ NTs obtained by SLM Ti6Al4V on gingival cells or oral bacteria. Xu et al used titanium grade II discs obtained by SLM, with a nanotube layer functionalized by calcium. After 7 days of culture, they showed improved proliferation and adhesion of gingival fibroblasts and keratinocytes on SLM surface with an Ra of approx. 2 µm [23]. Guo et al. proposed a protocol based on anodized grade V titanium surfaces obtained by SLM with an Ra of approx. 6 µm. They demonstrated that, at early stage (4h), hydrogenated NTs Ti6Al4V surfaces enhanced fibroblasts adhesion. Then, at day 1, their surface increased the expression of focal adhesion related genes (ITG-β1, FAK and VCL) [24]. These 2 studies deliberately combine a “micro and nano-topography” to enhance surface bioactivity but the high roughness may facilitate the bacterial adhesion in clinical conditions. Hu et al developed a nanotubes layer coated with CaP on grade II titanium surface obtained by SLM [25]. They observed that the elaborate surface significantly decreased the adherence of oral Streptococcus compared to native SLM surfaces, despite a Ra around 3 µm.

To the authors’ best knowledge, no studies combine a parallel assessment of procaryotes and eucaryotes’ behaviour on an SLM titanium surface with common NTs characteristics after anodization. However, this dual approach is crucial to better understanding the phenomenon that some refer as “race to invade”, where there is competition between host cell integration and bacterial colonization[6].

The formation of oral bacterial biofilms is a multifaceted, dynamic, and complex process that proceeds through several sequential stages involving diverse bacterial species [26]. Around natural teeth, the oral biofilms are initially predominantly colonized by Streptococcus species such as *S. gordonii*, which adhere to the tooth surface. These pioneering bacteria then facilitate the attachment of intermediary species like *Veillonella parvula* or *Fusobacterium nucleatum*, which serve as a bridge for the subsequent adhesion of late-colonizing bacteria, including opportunistic pathogens such as *Porphyromonas gingivalis* [27]. Peri-implantitis is characterized by even greater microbial diversity and a complex pathogenic biofilm bacteria an among these, *Porphyromonas, Treponema, Fusobacteria* and *Tannerella* have emerged as the main bacterial species associated with peri-implantitis [28–30].

Controlling the growth and adhesion of representative bacterial species on the titanium implant surface may effectively contain the cascade of events leading to biofilm maturation, thus reducing the risk of biofilm-related infections and biological complications of ISPs.

Furthermore, no study evaluates the bioactivity of SLM Ti6Al4V surfaces that have been pre-polished, following a dental laboratory protocol before the anodization. Yet, in clinical condition, the transgingival area of ISP is reworked technicians using handheld rotary burs to achieve the smoothest possible finish with the rules that have already been established regarding bacterial adhesion. This final polishing is often omitted for both the tested surfaces and the control ones, despite its clinical relevance.

Therefore, study aims to investigate how a polished anodized SLM Ti6Al4V surface, affects both the behaviour of Human Gingival Fibroblasts (HGFs) and a representative oral bacterium compared to the conventional titanium used today in ISP fabrication.

This parallel in vitro approach may provide clearer insight into the early events of implant colonization may help the development of mucosa-conscious implant surfaces.

## Material and methods

### 1) Surface manufacturing

Ti6Al4V discs with a diameter of 12 mm and a thickness of 2.5 mm were manufactured by an SLM machine (SLM 125 HL), with a Ti6Al4V ELI powder grade 23 (CT Powder Range Ti64 F, Carpenter Additive). SLM processing parameters are described in Table 1.

**Table 1:** SLM processing parameters used to manufacture Ti6Al4V SLM discs.

| **Laser power** | **Scanning velocity** | **Thickness layer** | **Hatch distance** | **Pattern** |
| --- | --- | --- | --- | --- |
| 275 W | 1100 mm/s | 30 μm | 120 μm | None |

Heat post-treatment was performed to achieve the mechanical properties required for dental prosthesis production [31]. The discs were heated in a furnace under an argon gas atmosphere, before a progressive heating was applied at a rate of 10-12 °C/min, up to 1050 °C for 2 hours. 1050°C/2 hours under vacuum followed by cooling in the furnace to 500°C and under argon gas below that temperature.

The polishing steps were first conducted on a rotational machine using P280, P320, P600, P800 and P1200 sandpaper. Then, all samples were manually polished (Fig. 1) with a specific dental bur (Dedeco® no. 4950, Long Eddy, NY) inserted into a dental handpiece with a slow speed ranging from 10.000 rpm to 12.000 rpm. Finally, a finishing bur (Robinson Polishing Bristle Brushes soft, Buffalo Dental Manufacturing Co Inc, Syosset, NY) combined with a universal polishing paste (Dialux Banc) was applied with the same conditions.

**Figure 1:**
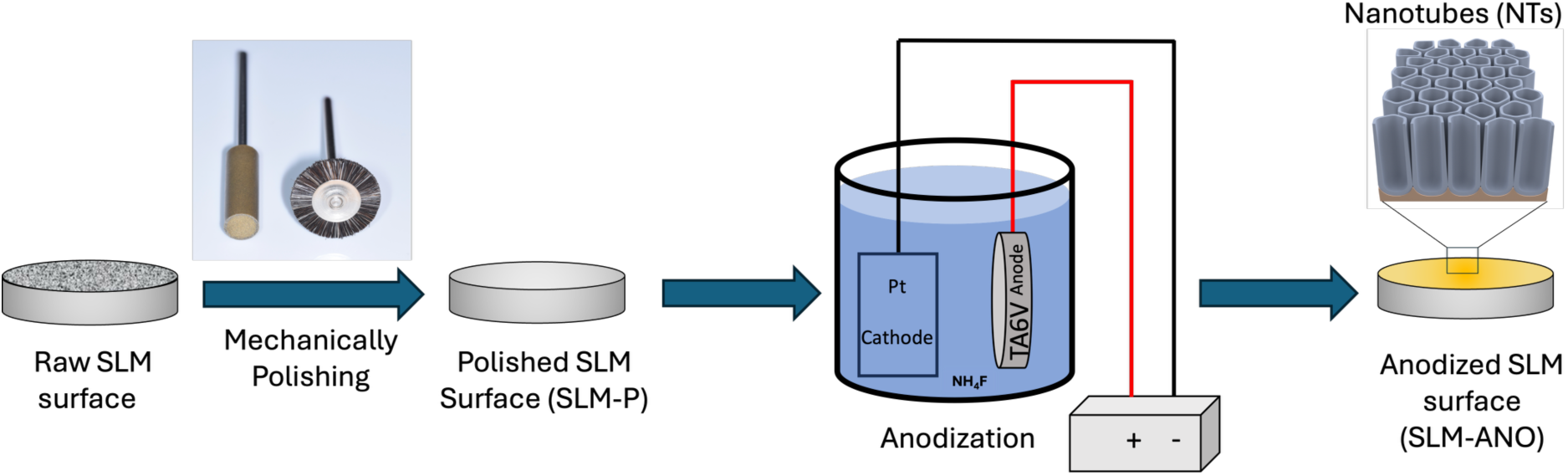
Schematic representation of the protocol to prepare the surface of interest (SLM-ANO).

The control discs were obtained using a conventional milling procedure from a Ti6AL4V grade 5 (Coprar Ti-5, Whitepeaks Dental Solutions GmbH, Germany). The same finishing burs were used to obtain the surface currently used in clinics.

### 2) Electrochemical Anodization

SLM polished discs (SLM-P) were washed into an ultrasonic bath of 75 wt.% ethanol and deionized water, before being anodized into a conventional two-electrode system (Fig. 1). The electrolyte solution was composed by 50 wt.% distilled water, 50 wt.% glycerol to increase viscosity and slow down ions transfer and 0.5 wt.% ammonium fluoride (NH_4_F) to supply fluoride ions. The discs were immersed into the electrolyte solution at room temperature (17-21 °C) for 50 min and submitted to an electrical voltage of 40 V. A platinum foil was used as a counter-electrode and placed approximately at 4-5 cm from titanium discs. The current intensity was recorded thanks to an amperemeter (EmjieBlue, Jeulin®). After anodization, the discs were rinsed with distilled water. The resulting surface of interest will be identified as “SLM-ANO” and the non-anodized control surface designated as “MP-CTRL”.

### 3) Surface characterization

#### 3.1) Surface topography

Nanoscale surface topography was observed thanks to a Scanning Electronic Microscope (SEM) with the accelerating voltage set at 15 kV (*Merlin, Carl Zeiss*®). Nanotubes’ length, pore diameter and wall thickness measurements were carried out on images obtained at low and high magnification.

#### 3.2) Roughness

Topographical analysis was performed using an 3D optical analyser (*Bruker Alicona®*) and MeasureSuite® software. The profile arithmetic average roughness (R_a_) and corresponding surface roughness parameter (S_a_) were evaluated on an 8 mm profile line and a 5 mm² surface area. Measurements were taken on three different zones, for 3 samples per condition. The average standard deviation (μm) of the data for of each group was calculated.

#### 3.3) Wettability

The water contact angle was measured using the sessile drop method in accordance with ISO 19403-2:2017. A 2 µL drop of ultrapure distilled water was deposited on the surface at room temperature. Its shape was captured by a camera after a few seconds of stabilization. The contact angle (CA) was calculated using the ellipse-fitting method. Measurements were carried out on 3 discs per condition on three different zones, with a drop shape analyser (*DSA25E Expert, Krüss®, Hamburg, Germany*) and its software (*Advance®*).

#### 3.4) Protein adsorption

Protein adsorption tests were performed with salivary proteins and Bovine Serum Albumin (BSA) coatings. For salivary proteins, saliva from four healthy and non-smoker donors was collected, and centrifugated at 9500 rpm for 40 min. The supernatant was collected and pasteurized at 60 °C for 30 min. Samples from the different donors were pooled and diluted into Phosphate Buffered Saline (PBS) (1:1) (pH7.4). Depending on the coating, a drop of 100 µL of this pasteurized saliva or BSA was incubated on anodized and control discs for 30 min, and specimens were rinsed three times in PBS. 100 µL of warmed (95°C) 2 % SDS (Sodium Dodecyl Sulfate) was pipetted onto the discs and incubated for 5 min. The SDS was then collected and diluted in PBS (1:20). Standards tubes and Micro BCA Working Reagent were prepared according to the Micro BCA Protein Assay Kit protocol (*ThermoFisher®*). In a 96-well micro-plate, 150 µL of each standard and sample were pipetted, and 150 µL of working Reagent was added. After an incubation for 2 hours at 37 °C, the absorbance of each sample and standards was read within 10 min using a spectrophotometer at 590 nm. Sample protein concentrations were obtained by comparison with the standard curve.

### 4) Eukaryotic cell biological assays

Fibroblast cells were obtained from human gingiva from the biological collection ORCELL (ethics approval n°19.11.04.64248 approved by the CER-SU on September 2023 (CER-2021-038) and expanded following an established protocol. Monolayer culture were maintained in fibroblast growth medium: Dulbecco’s modified Eagle’s medium (DMEM), low glucose, supplemented with 10 % foetal bovine serum (FBS) to promote cell growth, antibiotics (100 units/mL penicillin, and 100 μg/mL streptomycin), antifungal treatment (0.25 μg/mL amphotericin B), fibroblast growth factors (10 μg/mL βFGF) and 1 % nonessential amino acid, at 37 °C in a 5 % CO_2_-containing atmosphere. HGFs from three donors were cultured to consider inter-individual variability, aged between 23 and 26 years and used at a passage included between 3 and 5. As it is the surface currently used in clinics, polished milled discs (MP-CTRL) were used as the gold standard, and plastic slides with the same dimensions as discs were used as positive controls. For specimens’s decontamination, all discs were first washed in an ultrasonic bath of ethanol 75 % for 5 minutes and then exposed under UVC light for 30 minutes (*LK Sterilizer RTD-208A*).

#### 4.1) Cell proliferation and viability

Discs (SLM-ANO; MP-CTRL; Plastics) were place in a 24-well microplate. A cell suspension of 6 x 10^3^ cells/100 μL was prepared and an aliquot of 100 μL was added of the surface of each sample and cultivated at 37°C with 5% CO_2_ for 1 hour to allow cell attachment. Then, 900 μL of the medium was added into each well. At each time point (1, 4 and 7 days of culture) all the experiments were performed in triplicate for each 3 patients (total sample n=9 per condition). First, Live/Dead staining (*NucBlue Live Reagent/NucGreen Dead 488,* ThermoFisher®, *Waltham, Massachusetts, USA)* was used to evaluate cell viability. After 15 minutes of incubation, surfaces were observed thanks to a fluorescent microscope (*EVOS M5000, Invitrogen®*). For each disc, a count of living cells and total cells was performed using ImageJ in four random fields of view at a magnification of 20x for each experimental group. For cell proliferation evaluation, all wells were then trypsinized for 5 min at 37 °C and cells were counted twice using Malassez counting chambers.

#### 4.2) Cell resistance to enzymatic detachment

To determine adhesion strength against enzymatic detachment, protocol was inspired by Rivarii and al [32]. HGFs (from 1 donor) were seeded with an aliquot of 100 μL containing 5 x 10^4^ cells (n=4 specimens/condition), to cover the entire surface disc. After 6h of cell attachment, the discs were rinsed with PBS and incubated with 100 µL of 0,05 % trypsin solution diluted 1:10 in PBS, at room temperature for 20 min. At 1, 5 and 10 min of incubation, diluted trypsin solution containing detached cells was collected, and replaced. At 20 min, diluted trypsin solution was collected, and specimens were incubated into 100 µL of undiluted trypsin solution at 37 °C for 5 min, to collect the remaining adherent cells. In all tubes collected, cells were counted manually twice. Percentage of adhered cells was calculated by comparing the number of detached cells to the number of adherent cells. The same procedure was conducted after 36 hours of culture, with an initial cell concentration of 3 x 10^4^ cells per surface.

#### 4.3) Cell morphology

After 7 days of culture, HGFs were fixed with 4 % paraformaldehyde for 20 min (Formalin Sigma, Saint Louis, Mo, USA). Cells were then dehydrated using ethanol baths with increasing concentrations (50%, 60%, 70%, 85% and 100%). Samples were left in each bath for 5 min. After dehydration, a conductive palladium layer was sputted on the sample using a Cressington 208 HR sputter-coater monitored by a Cressington MTM 20 thickness controller. The specimens were then observed at hight magnifications, with the SEM celerating voltages between 5 and 15 kV.

#### 4.4) Immunofluorescence

In order to evaluate the expression of focal adhesions (FAs) in cell attachment induced by the surfaces, specific immunostaining for vinculin (VCL) was performed. Vinculin is a membrane-cytoskeletal protein that is involved in adhesion mechanisms. [33]. HGFs from one donor were seeded on different specimens at a density of 6 x10^3^ cells on each specimen (n=3 per condition) and cultured for 48 hours. Each disc was then fixed using a solution of 4 % paraformaldehyde and 5 % sucrose for 20 min and rinse 3 times. Then permeabilization with 0.5 % Triton for 10 min was performed and then rinses with PBS during 3 minutes containing 0.1 % Bovine Serum Albumin (BSA). A blocking solution composed of 1 % BSA and 0.1 % glycine was used to keep the pores open during 30 min. Primary antibodies were then immediately incubated overnight at 1/300. After three more rinses of three minutes with 0.1 % BSA, secondary antibodies adapted to the first antibodies were incubated for two hours, at a concentration of 1/400. The same rinsing protocol as the primary antibodies was applied, and Alexa Fluor® 594 phalloidin was used to stain actin on some specimens for 10 min. Finally, nuclei were marked with NucBlue (ReadyProbes, ThermoFisher®, Waltham, Massachsetts, USA) for 5 min. After being gently rinsed three times, images were then taken with a fluorescent microscope at x 4, x 10 and x 20 magnifications (EVOS M5000, ThermoFisher®, Waltham, Massachusetts, USA). Merged images were directly obtain from EVOS M5000 software.

Cell alignments were evaluated through immunofluorescence. After 7 days of culture, HGFs from on donor were seeded at 6 x 10^3^ per surface (n=3 per condition). Specimens were then fixed with a solution of 4 % paraformaldehyde and 5 % sucrose. After three rinses with PBS, cells were permeabilized with 0.5% Triton for 10 minutes. Cells were then incubated with phalloidin rhodamine X 1 for 15 minutes and DAPI for 5 minutes. Specimens were finally rinsed three times with PBS, before images were taken using a fluorescent microscope (EVOS M5000, ThermoFisher®, Waltham, Massachusetts, USA ®). Five images were taken for each specimen. Images were analysed using the Fiji ImageJ software, with the cell dispersion plugin.

#### 4.5) Gene expression

Cells were seeded at 20 x 10^3^ per surface (n = 4 per condition) and cultured for 48 hours. All specimens per condition were pooled and RNA genes were extracted (Tri-Reagent®Molecular Research Center, Euromedex) following the manufacturer’s instructions. RNA quantification was performed using the Qubit 2.0 fluorometer. cDNA was obtained by reverse transcription (RT), using the SuperScript II kit (ThermoFisher Scientific®) and following the manufacturer’s instructions. Real-time PCR (Polymerase Chain Reaction) was conducted using BIO-RAD CFX96 Touch (Bio-Rad Laboratories, Hercules, CA, USA). Diluted RT products were mixed with 7.5 μl of 2X Kapa SYBR Fast quantitative PCR (qPCR, Kapa Biosystems, Wilmington, MA, USA) and 300 nM primers for a final volume of 15 μl. Real-time PCR amplification was performed using the following program: one cycle at 94 °C for 3 min, 35 cycles at 95 °C for 5 s, gene-specific annealing for 20 s, and reaction completion with a reading plate and melting curve analysis from 65 °C to 95 °C, 5 s for each 0.5 °C. PCR reactions were performed in triplicate for each sample. Gene expression values were normalized with two reference genes (GAPDH and SDHA).

**Tableau 1:**
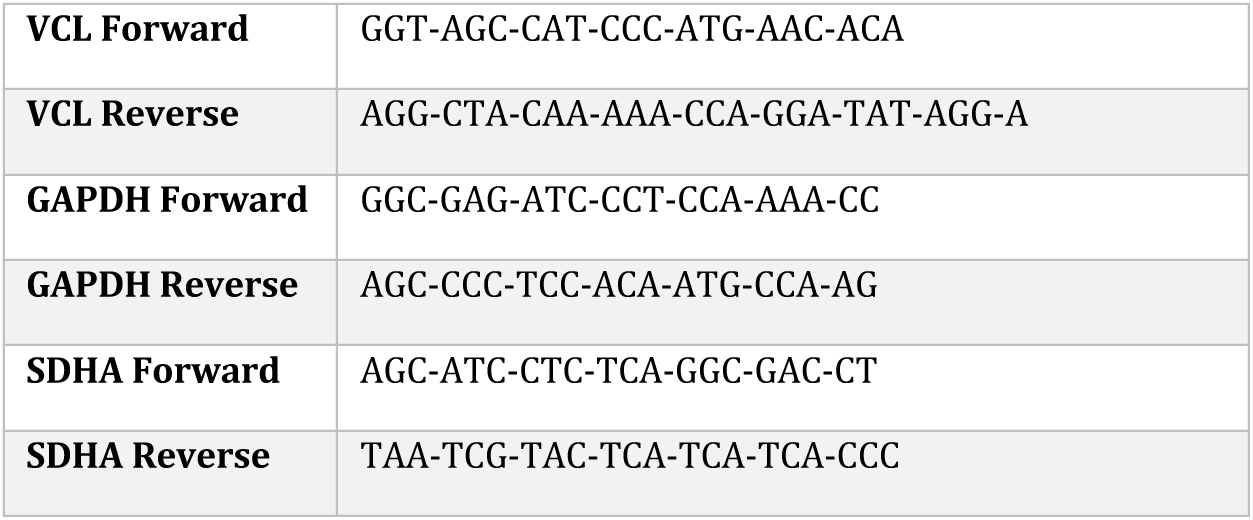
Target cDNA primer sequences used in quantitative polymerase chain reaction. VCL Vinculine; GAPDH glyceraldehyde 3-phosphate dehydrogenase; SDHA succinate dehydrogenase complex flavoprotein subunit A.

### 5) Bacteriological culture

#### 5.1) Titanium disk preparation

Polished Ti disks (n=6) and Selective Laser Melting (SLM) anodized titanium disks (n=6) were used in this study. Polished disks are used as control and had a diameter of 12.0 mm and thickness of 3.0 mm. SLM titanium disks had a diameter of 12.0 mm and thickness of 2.5 mm. Each disk was preliminarily sterilized in an ethanol bath for 10 minutes, followed by exposure to UV light for 30 minutes on each side.

#### 5.2) Bacterial culture and biofilm formation

*Streptoccoccus gordonii* strain DL1 was grown overnight in Brain Heart Infusion (BHI) agar and then inoculated into 5mL of BHI broth overnight at 37°C under anaerobic conditions to obtain bacteria suspension. One-quarter of the overnight culture was used the following day, to prepare a fresh bacterial suspension. The optical density (OD) was measured and adjusted to 0.2 at 600 nm, then transferred into a fresh BHI tube and incubated under the same anaerobic conditions for 3–4 h. Once the OD reached the desired range (0.3–0.5), the suspension was diluted to obtain a final OD of 0.05 in 3 mL of BHI broth for subsequent disk inoculation. Disks were incubated in 24-well plates and incubated with 1 mL of S. gordonii suspension (OD = 0.05 in BHI) under anaerobic conditions at 37 °C for 30 min to allow bacterial adhesion. After this pre-incubation, disks were transferred into fresh wells containing 1 mL of sterile BHI broth and further incubated anaerobically at 37 °C for 24 h to allow biofilm development.

#### 5.3) Biofilm detachment by sonication

After 24 h incubation, each disk was gently rinsed once with BHI to remove planktonic cells and transferred to a 15 mL Falcon tube containing 2 mL of BHI broth. Tubes were vortexed for 2 min, followed by sonication for 10 min in an ultrasonic bath (Branson 5800, Branson Ultrasonics, Danbury, CT, USA) to dislodge biofilm cells. A second vortexing step of 2 min was then performed to ensure complete detachment.

#### 5.4) Viability assessment

Serial ten-fold dilutions (up to 10⁷) of the detached bacterial suspension were prepared in a 96-well plate by mixing 180 µL of BHI broth with 20 µL of the sample. Triplicate aliquots from each dilution were plated on BHI agar and incubated aerobically at 37 °C with 5 vol.% CO₂ for 24 h. Colony forming units (CFU) were counted, and viable cell numbers were determined.

### 6) Statistical analysis

Data analysis was carried out using Prism software (GraphPad v.10.3.1). Kruskall-Wallis tests were used to compare roughness parameters; wettability; salivary proteins adsorption and gene expression results. BSA proteins’ adsorption data were treated with an unpaired T-test. For cell proliferation, cell viability and cell alignment, an (2-way or 1-way) ANOVA associated with Tukey’s multiple comparisons test were perform since the data were distributed according to the normality. Mann-Withney test were performed for analyse bacterial results. A p-value < 0.05 was considered significant.

## Results

### 1) Surface characterization

SEM images of the Ti6Al4V samples after anodization showed a nanotubular surface, with nanotubes exhibiting a diameter of around 100 nm and a wall thickness of 15 nm (Fig. 2). To measure their length, a deliberate scratch was made by a thin blade to observe the NTs profile. The nanotube length was estimated at 500 nm.

**Figure 2:**
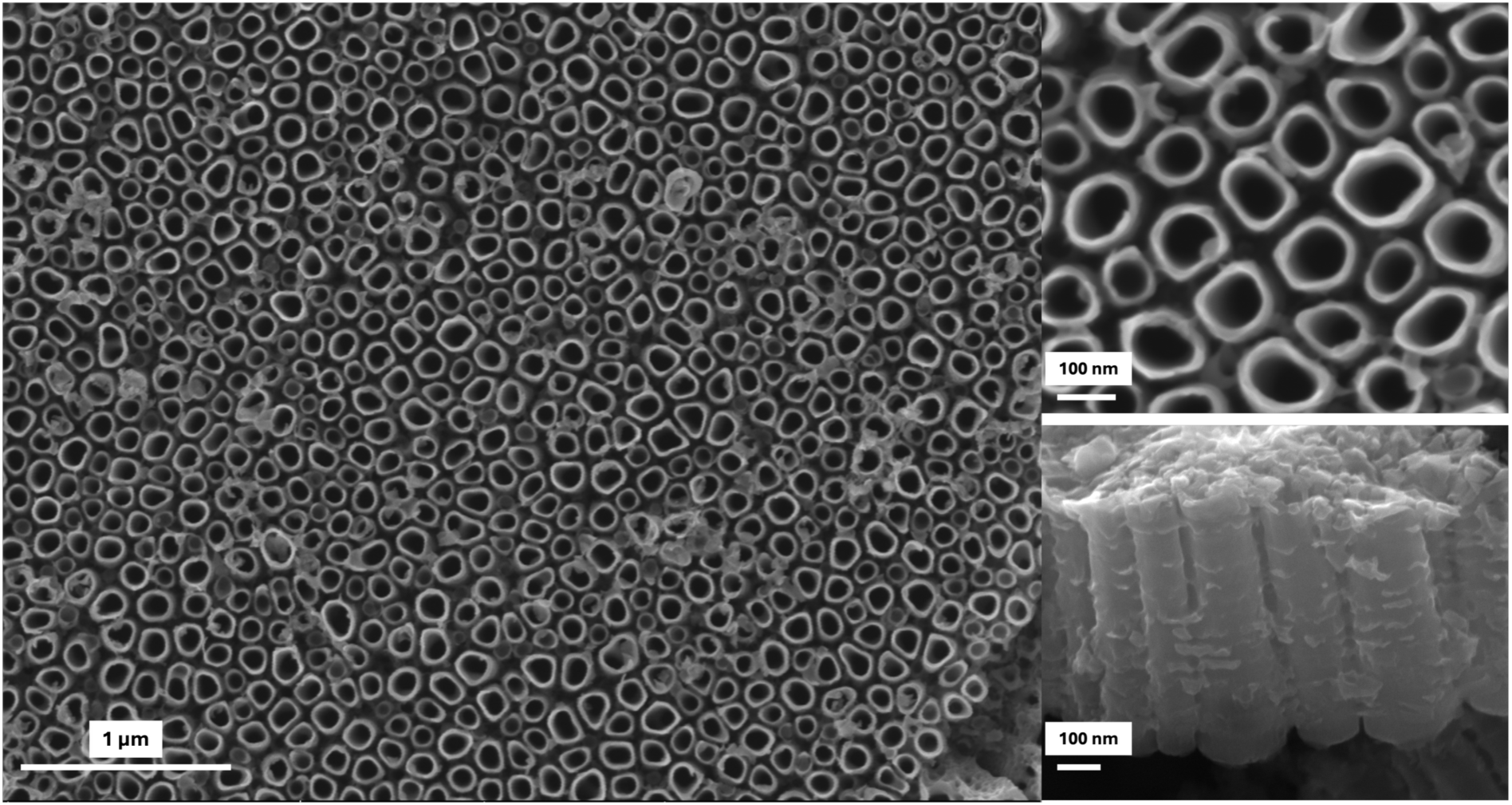
SEM images of NTs on a top view (x20.000 and x100.000 respectively) and a profile view (x100.000). Nanotubes show a diameter of around 100 nm, a wall thickness of 15 nm and a length estimated at 500 nm.

This nanostructuration does not appear to modify the micrometric roughness (Fig. 3a): the arithmetic average roughness (Ra) was similar between the SLM anodized discs (SLM-ANO) with a Ra equal to 0.198 µm ± 0.001 and the SLM polished discs (SLM-P) (Ra = 0.223 µm ± 0.001). However, a slightly statistical difference was notable between SLM-ANO and MP-CTRL (the machined polished control) with a Ra equal to 0.249 µm ± 0.003 (p<0.05). Furthermore, no statistical difference was shown in Sa values between these surfaces (p>0.05) with Sa= 0.24 µm ± 0.05; 0.25 µm ± 0.8; 0.29 µm ± 0;01 for SLM-ANO; SLM-P and MP-CTRL respectively. About wettability, contact angle (CA) measurements demonstrated that hydrophilicity increased for SLM-ANO discs (Fig. 3b). The average distilled water CA was significantly different between SLM-ANO (29.5° ±7.7) and SLM-P (73.4° ±5.9) (p<0.0001) and SLM ANO and MP-CTRL (63.3° ±5.5) (<0.005). Protein adsorption assays showed no statistical differences (p>0.05) between both surfaces, for salivary proteins and also for Bovin Serum Albumin (BSA) (Fig. 3c).

**Figure 3:**
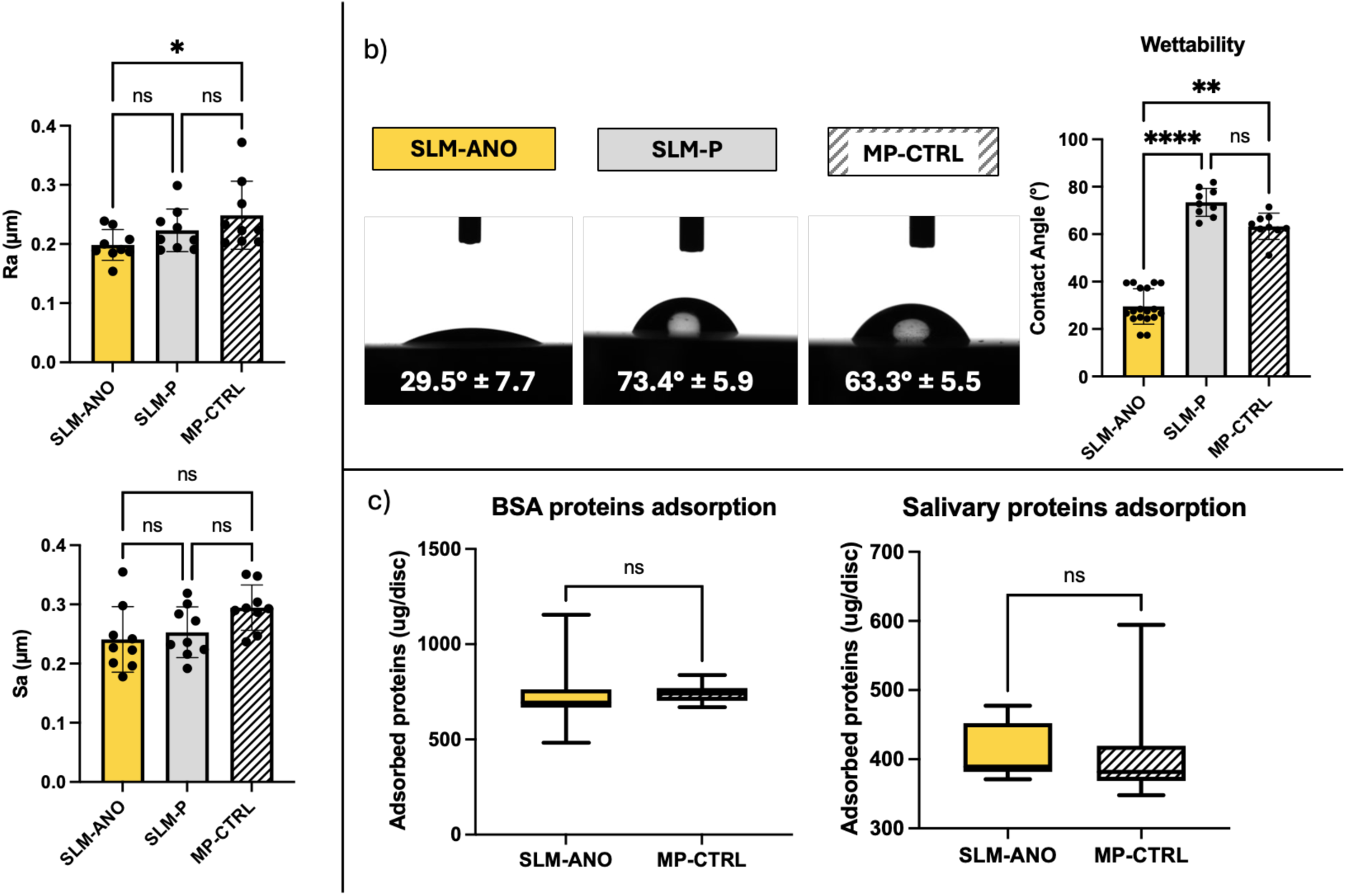
a) Average roughness (Ra) and surface average roughness (Sa) * (p<0.05) b) Contact angle measurements **** (p<0.0001); ** (p<0.005) c) BSA proteins adsorption and salivary protein adsorption on Ti substrate.

### 2) Eukaryotic cell biological results

#### 2.1) Cytocompatibility

Regarding cell proliferation and viability (Fig. 4a), there were no statistical differences within the groups after 1, 4 and 7 days of culture (p>0.05). Proliferative activity was particularly marked between D1 and D4 with a p<0.001 for all group, with a stagnation between D4 and D7 for all group (p>0.05). Cell viability was above 95% on all surfaces and at each time point.

**Figure 4:**
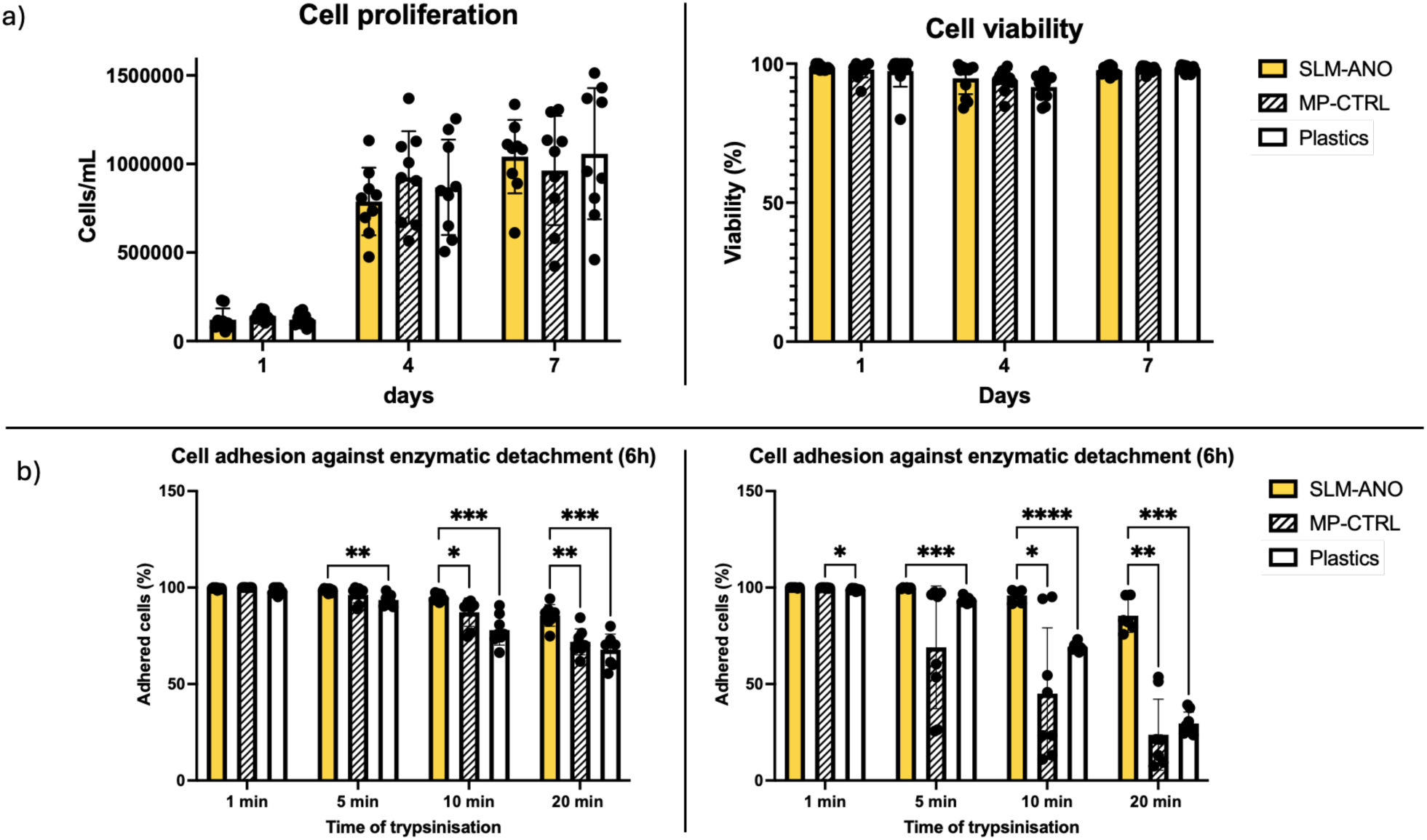
a) Cell proliferation and cell viability at 1, 4 and 7 days on the tested surface (SLM-ANO), the control surface (MP-CTRL) and the positive control surface (Plastics). There was no statistically significant difference between the different groups at all time point (p>0.05). b) Cell adhesion against enzymatic detachment at 6 and 36 hours.**** (p<0.0001); *** (p<0.001); ** (p<0.01); * (p<0.05). After 10 and 20 minutes of trypsinization, the percentage of adherent cells was higher on anodized surfaces than on other groups, at short term (6 h) (p<0.01) and middle term (36 h) (p<0.01)

#### 2.2) Resistance to enzymatic detachment

HGFs were more resistant to trypsinization on SLM-ANO discs, than on MP-CTRL and plastic surfaces at short term (6h) and middle term (36h) (Fig. 4b). After 6h of culture to allow attachment, following with 20 min of trypsination, the number of adhered cells was higher on SLM-ANO (85.5 % ± 5.6) than MP-CTRL and Plastics (71.9 % ± 6.7 and 87.8 % ± 8.1). At 36 h of culture, the difference was significant after 10 minutes of trypsinization with 95.9 % ± 3.2 of adhered cells compared to 45.0 %± 34.11 (p<0.01) on MP-CTRL and 69.0 % ± 2.0 on Plastics (p<0.001). Interestingly, the percentage of adherent cells was above 50 % on all surfaces after 6 h of adhesion, but at 36 h of culture, this percentage dropped to less than 25 % after 20 min of trypsinization on the MP-CTRL. On SLM-ANO samples, this percentage remained above 75 % at all time, regardless of the duration of the culture.

#### 2.3) Cell morphology, immunofluorescence and cell alignment

SEM images obtained after 7 days of culture exhibited the presence of numerous filopodia on SLM-ANO surfaces, structures used by the cells to adhere to the substrate. Figure 5 shows an example of filopodia on the nanotube layer at a very hight magnification.

**Figure 5:**
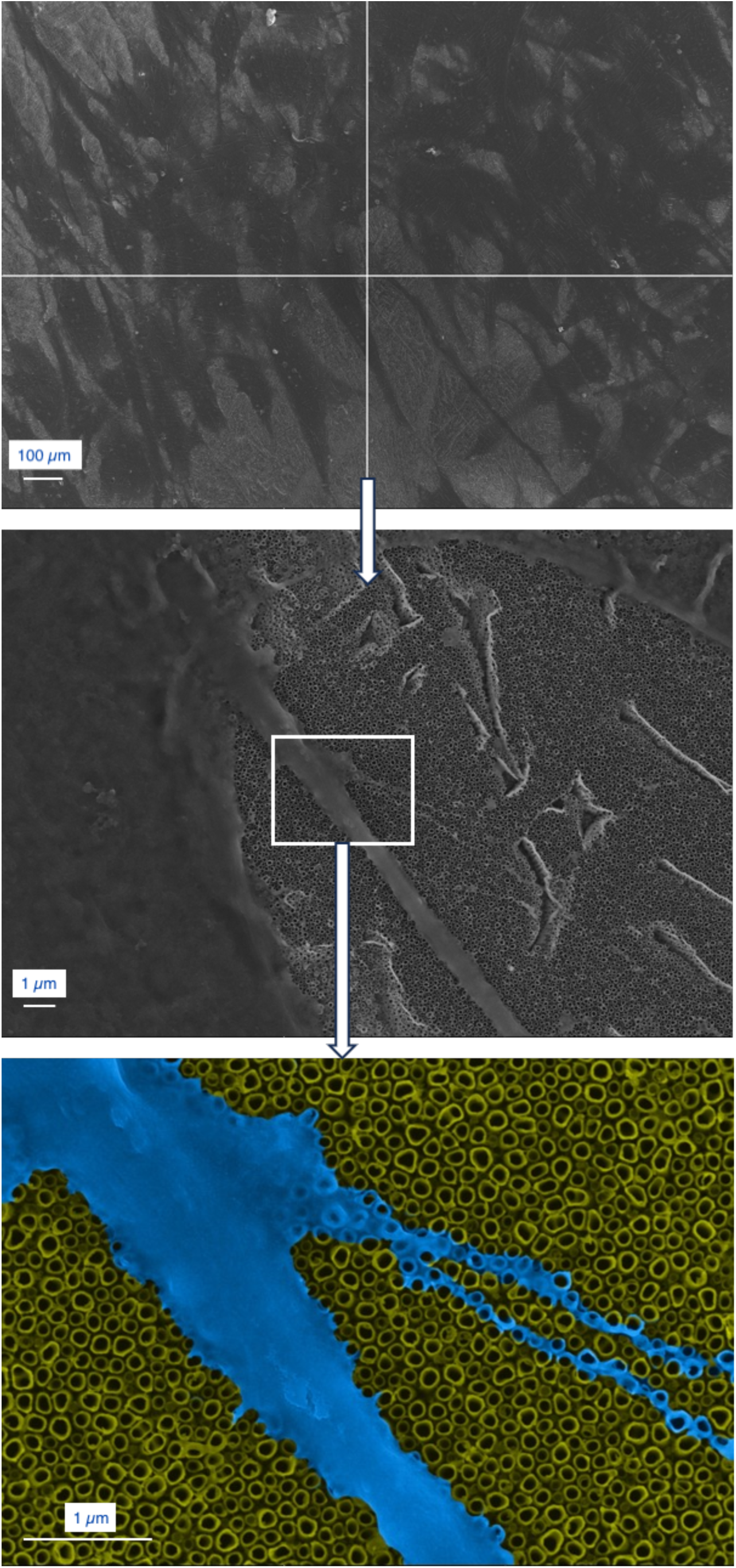
SEM images of HGFs cultured on Anodized surfaces (SLM-ANO) at 7 days. Magnification x100: x5.000; x100.000 of the same area. On hight magnification, it can be observed that filopodia (which emanate from lamellipodia) anchors to the edges of the NTs)

On SLM-ANO, vinculin VCL was homogeneously expressed into the cell membrane. In contrast, on the MP-CTRL samples, the distribution was located preferentially located at the edges of the cells (Fig. 6a). Regarding cell alignment, the similarity between the MP-CTRL samples and the surface of interest SLM-ANO was demonstrated by the fact that the cells spread out in a uniform direction, compared with plastic surfaces where the cells spread out without any preferential axis (Fig. 6b).

**Figure 6:**
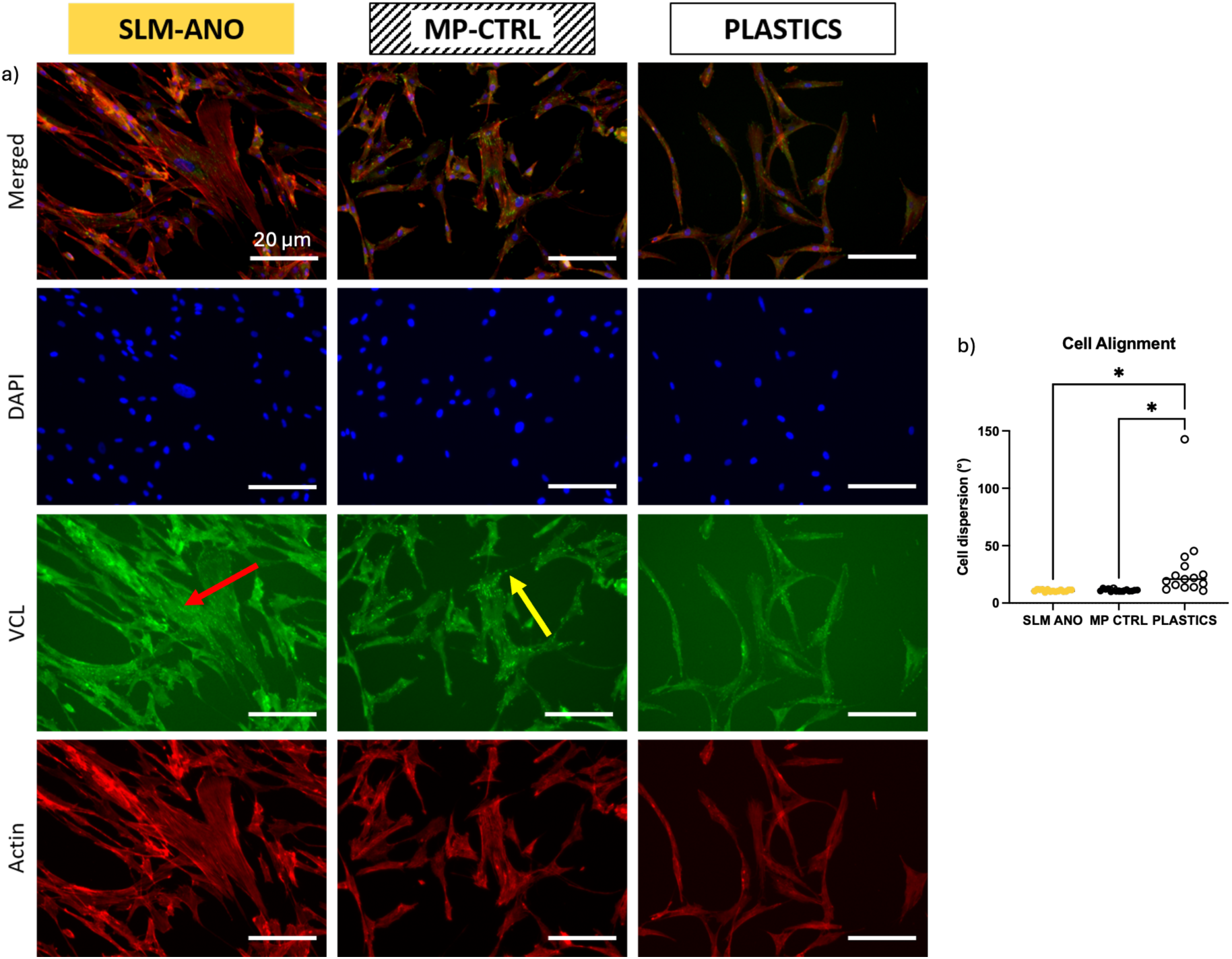
a) Cytoskeletal organisation analysis with immunofluorescence staining. The HGFs were cultured on the SLM-ANO surfaces, MP-CTRL surfaces and Plastics for 48 h. The cells were labelled for vinculin (green), F-actin (red) and nuclei (blue). On the controlled surfaces, vinculin signal was detectable at the edges of the cells, whereas on the anodized surfaces (yellow arrow), the signal was visible more uniformly (red arrow). b) Cell alignment after 7 days of culture. On the tested surfaces (SLM-ANO) and control surfaces (MP-CTRL), cell distribution was statistically more uniform (p<0.05) than on plastic surfaces, where cells proliferated without any particular orientation.

#### 2.4) Gene expression

The presence of nanotubes regulated vinculin (VCL) gene expression of HGFs after 48 hours of culture. VCL was upregulated on the surface of interest (SLM-ANO) compared with the control surface (MP-CTRL) and the plastics surfaces (p<0.05) (Fig. 7).

**Figure 7:**
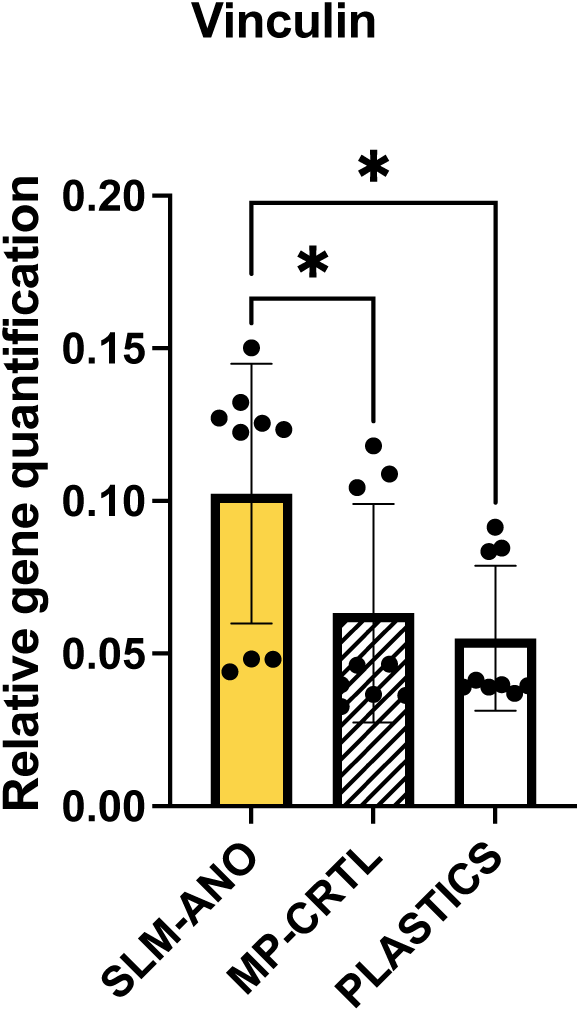
Gene expression of vinculin. The SLM-ANO surface exhibiting the presence of nanotubes increased the expression of vinculin at 48 h compared to the control surface and the plastics (* p<0.05).

### 3) Bacteriological results

To assess the capacity of the newly developed surface to control bacterial adhesion, we estimated the capacity of the early surface colonizer *Streptococcus gordonii* to colonize SLM-ANO vs MP-CTRL titanium disk. The colonization capacity of *Streptococcus gordonii* was equivalent on the SLM-ANO and control surfaces (MP-CTRL) with a level of colonization around 105 CFU/mm^2^ (p= 0.7052) (Fig. 8).

**Figure 8:**
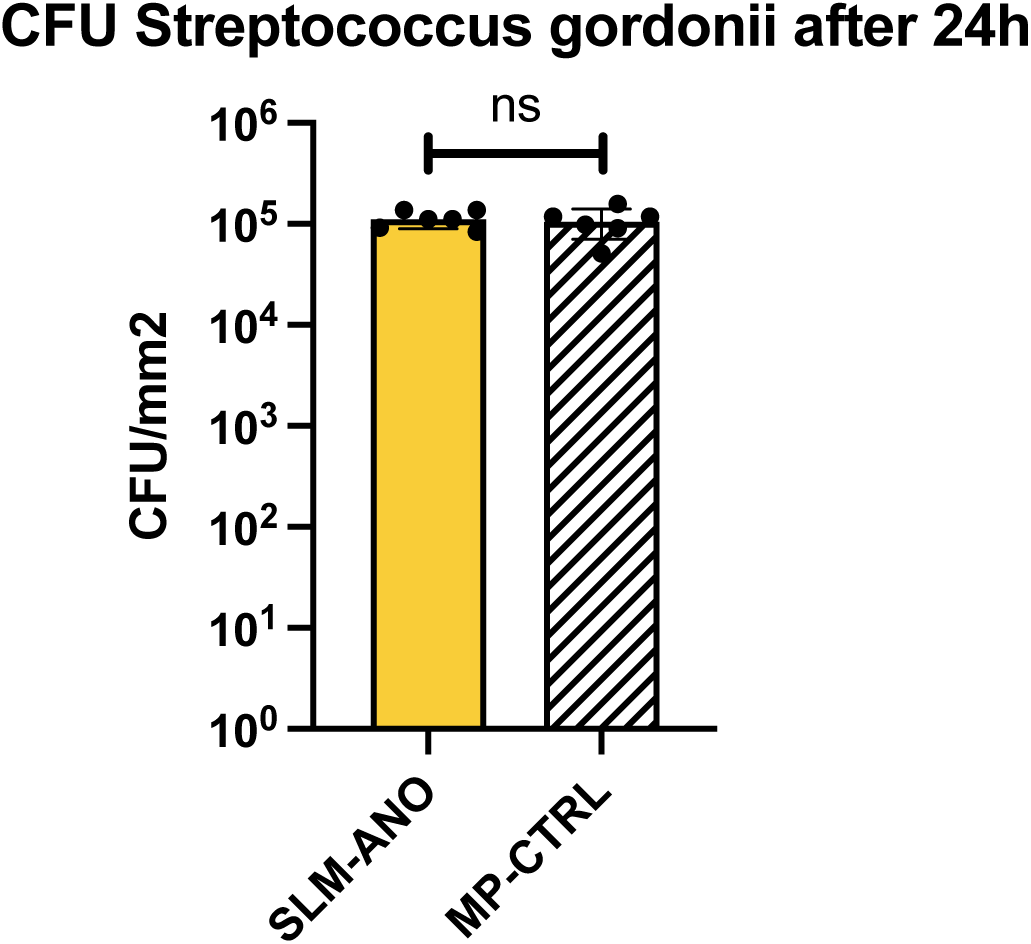
Biofilm formation of Streptococcus gordonii on titanium disks. Sterilized titanium disks were incubated with 10^5^ cells of S. gordonii for 10 min to allow for initial adhesion and transferred into fresh BHI medium for 24h at 37°C under anaerobic conditions to allow biofilm development. Biofilm cells were dislodged from the disks by vortexing and sonication and colony forming units were determined. They were no significant difference between SLM-NO and MP-CTRL (p>0.05): the SLM-ANO surface does not appear to be as readily colonized by this bacterium.

## Discussion

This study demonstrates that anodized nanotubular surfaces fabricated on SLM Ti6Al4V alloy, using a clinically relevant polishing protocol, improve human gingival fibroblast adhesion, resistance to enzyme detachment, without promoting *S Gordonii* colonization.

Peri-implant diseases remain a major health issue and surface modifications at a nanometric scale are a promising solution to improve soft tissue integration. Here we chose anodization to only nanostructure the titanium surfaces, since it is reproductible, and relatively easy to perform. Anodization can be used as pre-treatment for metallic nanoparticles grafting or antibiotic filling. However, some concerns exist regarding the long term safety of metallic nanoparticles and the effectiveness of antibiotic loading [34]. In addition, we decided to apply a protocol feasible at chairside.

For this study, Ti6Al4V discs were obtained by SLM. As for the MP-CTRL control surface, a mechanical polishing protocol was applied, to fit with the methods currently used to manufacture ISP transgingival parts in prosthetic laboratories. The anodized protocol enabled us to obtain organized nanotubes on the SLM surfaces, presenting an average diameter of 100 nm, a 15 nm wall thickness and 700 nm length (Fig. 2).

These nanotube dimensions are consistent with previous results, since gingival fibroblasts exhibit a better adhesion on NTs presenting a diameter between 70 and 100 nm [35,36]. Furthermore filopodia developed by cells are able to detect patterns above 15 nm [37], which is the NTs wall thickness that were obtained. Regarding surface characteristics, anodization does not affect micrometric roughness since Ra values are comparable to the surface before modification (Fig. 3a). This roughness around 0.2 µm is compatible with the threshold approved by consensus [38]. Concerning surface biocompatibility, HGFs proliferated similarly on the SLM-ANO surfaces that on the control surfaces (Fig. 4a). Cell viability was also similar with a viability exceeding 95 % for both groups. Cell proliferation and viability were assessed considering inter-individual variability by using cells from different human donors. Interestingly, eukaryotic cell adhesion results reported that HGFs cultivated on SLM-ANO surfaces showed a better resistance to enzymatic detachment than HGFs cultivated on control surfaces (Fig. 4b). These, short-term (6 h) and middle-term (36 h) results were correlated with cell morphology at 7 days, where HGFs showed filopodia firmly anchored on NTs edges (Fig. 5). Many other studies reported the presence of these adhesive structures on anodized surfaces [16,33] but as far as we know, very few very high magnification images are currently available.

The uniform distribution of vinculin, throughout the cell membrane on SLM-ANO could explain why adhesion was better on SLM-ANO compared to MP-CTRL where vinculin was only located at the cell’s outer edge (Fig. 6). Furthermore, vinculin was upregulated by HGFs cultured in contact with nanotubular SLM surfaces (SLM-ANO) (Fig. 7). In other words, SLM-ANO surfaces appear to promote both, vinculin production and distribution within the cell. Nevertheless, this improvement is probably due to the NTs and not to the surface manufacturing method, since other teams have reported similar results on nanotubular titanium surfaces obtained by machining. Wang et al. [33] showed that HGFs in contact with Ti surface containing NTs of 100 nm, upregulated the gene expression of VCL at 24 h. Similarly, at 7 days, Xu Z and al. [39] also reported a higher expression of VCL of HGFs cultured on NTs with a diameter of 100 nm.

The common spreading and alignment of HGFs on anodized SLM and control surfaces could be explained by the polishing protocol applied before anodization, which can leave polishing grooves on a microscopic scale. These preliminary grooves can guide the formation of NTs and help to establish a regular pattern, as described by Gulati [40,41]. Cell alignment plays an important role in EMC remodeling and cytoskeleton organization [42].

Hydrophilicity was increased for anodized SLM surfaces. It has been reported that hydrophilic surfaces interact closely with biological fluids such as blood or saliva, allowing normal protein adsorption to the surface [^43^], which is the first step when implanting a biomaterial into the human body. Then, these proteins are then used to anchor cell receptors to adhere to the surface and trigger a cascade of intra-cellular reactions leading to the production of the ECM. Despite improved hydrophilicity, the adsorption of salivary proteins or BSA remained unchanged. These result are comparable with Xu Z and al. [39] who reported that that protein aggregates (30 nm size regime) are too small to anchor on NTs with a diameter of 100 nm. However, they reported that fibronectin (120 nm size) could be more easily adsorbed onto these same NTs. Similarly, Li et al highlighted an increased protein adsorption with a higher nanotubes diameter (>180nm) [44] but they also indicated that a wide-open NTs disconnect the ECM and could damage the development of focal adhesion and therefore recommends NTs with a small diameter.

With regard to the outcomes of bacterial colonization analysis, anodization did not manifest as either a positive or negative influence on the biofilm formation capacity of the early colonizer S. gordonii. The bacterial proliferation was similar that on the control surface currently used in practice, which meets the minimum requirement for clinical translation (Fig. 8). This is relatively consistent with results from the literature that generally detected either no or a moderate (< 1 log in CFUs) impact of titanium anodization on the adhesion of various bacterial species [25,39,45]. Nevertheless, a comparison of different studies is challenging due to variations in the anodization process and the resulting properties of the formed nanotubes [46]. In addition to the morphology of the nanotubes, a recent systematic review and meta-analysis reports that the antibacterial effect of nanotubular surfaces alone varies depending on the bacterial strain used. They only demonstrated significant bacterial reduction in studies investigating *Staphylococcus aureus* over a period of 6 hours [47]. Then, a 24-hour experiment may erase the effect of the nanotubes on bacteria.

A further interesting and not negligible point concerns the color of the discs that it was possible to obtain using this protocol. After anodization, the discs were yellow/gold, which was reported to be an aesthetic advantage for masking the titanium abutment underneath the gingiva [48].

It should be noted that these promising results were obtained from human gingival fibroblast (HGFs) located in the connective part of the peri implant soft tissues. However, these tissues comprised two major structural cell types, fibroblasts and epithelial cells, with different embryonic origins and different behaviors [15]. Therefore, additional investigations with human epithelial cells will be required to supplement the presented results and confirm the biocompatibility and the clinical relevance of the SLM-ANO. In the meantime, the bacterial evaluation focused on single-species bacterial cultures, and it is obvious that the oral ecosystems are much more complex structures that cannot be explained so straightforwardly.

## Conclusion

The nanotubular SLM titanium alloy surface, easy to obtain and reproductible appears to promote the in vitro adhesion of human gingival fibroblasts, while not enhancing bacterial adhesion and proliferation of an oral early colonizer. Further investigations with epithelial cells and multispecies bacteria are needed to better simulate clinical conditions.

## Acknowledgements

The authors are grateful to Edentech for providing specimens. The authors would like to thank Nil Bouaouni for the 3D illustration and Bruno Stauder from Bodycote for heat treatment.

